# Merkel cell polyomavirus in Merkel cell carcinoma: Integration sites and involvement of the KMT2D tumor suppressor gene

**DOI:** 10.1101/2020.08.03.234799

**Authors:** Reety Arora, Jae Eun Choi, Paul W. Harms, Pratik Chandrani

## Abstract

Merkel cell carcinoma (MCC) is an uncommon, lethal cancer of the skin caused by either Merkel cell polyomavirus (MCV) or UV-linked mutations. MCV is found integrated into MCC tumor genomes, accompanied by truncation mutations that render the MCV large T antigen replication incompetent. We used the open access HPV Detector/ Cancervirus Detector tool to determine the MCV integration sites in whole exome sequencing data from 5 MCC cases, thereby adding to the limited published MCV integration site junction data. We also systematically reviewed published data on integration for MCV in the human genome, presenting a collation of 123 MCC cases and their linked chromosomal sites. We confirm that there are no highly recurrent specific sites of integration. We found that, chromosome 5 is the chromosome most frequently involved by MCV integration and that integration sites are significantly enriched for genes with binding sites for oncogenic transcription factors such as LEF1 and ZEB1, suggesting the possibility of increased open chromatin in these gene sets. Additionally, in one case we found integration involving the tumor suppressor gene *KMT2D* for the first time, adding to previous reports of rare MCV integration into host tumor suppressor genes in MCC.

## INTRODUCTION

Merkel cell carcinoma (MCC) is an aggressive skin cancer (5 year overall survival rate of about 44% and disease-specific mortality of 33-46% ^1,2^), predominantly arising in the elderly and the immunocompromised, that is associated with Merkel cell polyomavirus (MCV) in the majority of cases^2,3^. MCV was discovered in 2008 by Feng et al., and was found to be integrated into the MCC tumor genome in a mutated, replication-deficient form^3^. MCV is a small (5 kb), non-enveloped, circular, dsDNA polyomavirus that is commonly found on human skin^2–4^. The MCV early region expresses large T antigen (LT) and small T antigen (sT) proteins, that have been shown to drive tumorigenesis and are likely causative for MCC^5,6^. MCC tumor cells express a truncated form of LT protein that cannot mediate viral replication, but retains the domain responsible for inhibition of the tumor suppressor Rb^2–5^. MCC that lack MCV display cellular genomic mutations in tumor suppressor genes, especially *TP53* and *RB1*^*7–9*^. Additional tumor suppressors such as *NOTCH* family genes and *KMT2D* are inactivated at lower rates in MCV-negative MCC^2,7–9^.

Integration into the human genome is implicated as an early oncogenic event for many DNA tumor viruses^10,11^. Integration increases the risk of cancer beyond simple viral infection as it can disrupt and deregulate nearby human cancer-associated genes, contribute to genome instability, and create fusion transcripts^11,12^. Various studies show an overall bias for integration near open chromatin regions and SINE elements for DNA tumor viruses^12^. However, interestingly there are differences between integration site patterns related to different virus types, cancer types and disease stages.

Because MCV infection is near-ubiquitous^10^, demonstration of viral integration can be helpful in distinguishing tumor-associated viral sequences from spurious detection of background virus. Furthermore, viral integration can interrupt tumor suppressor genes, hence representing a potential mechanism for viral tumorigenesis in addition to viral oncogenes. Despite the diagnostic and biologic importance of this question, studies investigating MCV integration sites have been relatively few, due to the rarity of MCC and limited high-quality tumor samples. Furthermore, the identification of MCV integration is technically challenging, as the circular genome of the virus does not linearize in a consistent manner. Several approaches have been developed to address this question, often relying upon custom assays or analysis tools (Table 1 and Supplementary Table 1). A sensitive approach for detecting MCV integration sites using commercially available assays and open-access analysis tools has not yet been uniformly validated and adopted.

**Table 1:** Summary of Studies on MCV Integration in the Human Genome.

| Year | First Author (Journal) | Detection Assay | Integration Analysis Tool | Number of MCC - Integration Sites Detected | Human Genome Integration Junction, Chromosome (# of cases) | MCV Genome Integration Junction (# of cases) |
| --- | --- | --- | --- | --- | --- | --- |
| 2008 | Feng H. (Science) | cDNA pyrosequencing, 3'-RACE, 5'-RACE, Southern Hybridization | Digital Transcriptome Subtraction | 2 | chr3 | LT (1) |
| 2009 | Sastre-Garau X. (J Pathol) | DIPS-PCR | N/A | 10 | chr2, 3, 4, 5, 6, 8, 12, 20, and Y | LT (7), NCCR (2), VP1 (2) |
| 2010 | Laude H.C. (PLoS Pathogens) | DIPS-PCR | N/A | 4 | chr6, 9, 11, 15 | LT (3), VP1 (1) |
| 2011 | Duncavage E.J. (J Mol Diagn) | MCPyV-specific Hybrid Capture NGS | SLOPE and Breakdancer | 4 | chr6, 8, 9 | Not reported |
| 2012 | Martel-Jantin C. (Virology) | DIPS-PCR | N/A | 19 | chr1 (3),3 (1),4 (3), 5 (4), 6. (2), 14 (1), 18 (1) and 19 (2) | LT (13), ST(1), VP1(3), VP2(2) |
| 2013 | Guastafierro A. (J Virol Methods) | Lambda phage library ,3'RACE, 5'RACE and Southern hybridization Library, | N/A | 4 | chr3, 5, 6, 10 | LT (3), ST (1) |
| 2015 | Doolittle-Hall et al., Cancers | N/A(Meta-analysis of published reports) | Meta-analysis of published reports | 37 | Multiple, Non-recurrent | Not reported |
| 2017 | Starrett G.J. (mBio) | Whole Genome NGS | Custom Pipeline | 4 | chr1, 6, 19, 20 | LT (2), NCCR (2) |
| 2019 | Schrama D. et al., (Int J Cancer) | DIPS-PCR | N/A | 5 | chrs 1,2,6,11,12 | LT (2), VP1 (4), VP2( 3) |
| 2020 | Slevin M.K. (J Mol Diagn) | Hybird-capture and Massively parallel sequencing (Viropanel) | SvABA (structural variation and insertion/deletion analysis by assembly) | 33 | chr1 (5), 2 (2), 3 (1), 5 (7), 6 (4), 7 (2), 8 (2), 9 (2), 10 (1), 11 (4), 15 (1) and 16 (2) | LT (5), ST (2), VP1 (3), VP2 (1) and **(1) |
| 2020 (bioRxiv) | Czech-Sioli M. (bioRxiv) | Long-read third generation NGS, Nanochannel and Nanopore sequencing | Multiple | 9 | chr3, 4, 5 (4), 9, 11,13,20 | LT (5), NCCR (2), VP1 (8), VP2 (4) and ** (1) |
| 2020 (bioRxiv) | Garcia-Mulero S. (bioRxiv) | Human coding region sequencing NGS (Hybird Capture) | No specialized analysis tool | 1 | chr 19 | Not reported |
| 2020 | Starrett G.J. (Genome Med) | Hybird-capture and Massively parallel sequencing (Viropanel) | Custom pipeline ( oncovirus tools- suite - (https://github.com/gstarrett/ncovirus_tool s)) | 25 | chr1 (4), 2(2), 3(1), 5(4), 6(3), 7(2), 8(2), 9(3), 10(1), 11(1), 15(1), 16(2) and 18(1) | Not reported |
| 2020 | Arora R. (current study) | whole exome (Hybrid Capture) | HPVDetector | 5 | chr4, 7, 12, 16 and 17 | LT, NCCR (2), VP1 (2) |

\*\* This was identified in the region between MCV LT and VP1 coding regions in the MCV genome

MCV integration sites have been reported across most chromosomes, and may occur within genes or in intergenic regions (Table 2). With rare exceptions, the integration site for a particular tumor (and its related metastases) is unique to that case. Despite this variability, several studies have suggested that the distribution of integration sites is non-random, with patterns related to mechanisms of integration^8,9^. Unlike HIV, MCV does not carry an integrase, and integration into the host genome is not a part of viral life cycle^3^. Starrett et. al. reported that integration is likely mediated through erroneous DNA repair at sites of microhomology between the host and viral genomes, similar to mechanisms identified for HPV genome integration in squamous cell carcinoma tumors^8,9^. Similarly, Czech-Sioli et al. recently observed that MCV follows two main types of integration site structure: (1) linear patterns with chromosomal breakpoints related to non-homologous end joining, and (2) complex integration loci dependent on microhomology-mediated break-induced replication^13,14^. Doolittle-Hall et al., the first meta-analysis done on 37 MCCs for integration analysis, found that MCV integration occurred preferentially near SINEs and BDP1 binding sites^12^. They also confirmed viral preference for transcriptionally-active gene-dense regions and accessible chromatin. In addition, they observed a tropism toward integration near specific categories of genes: sensory perception and G-protein coupled receptor genes^12^. MCV integration within cellular genes could also act as a mechanism of tumor suppressor gene inactivation in a subset of tumors, as potentially disruptive integration into genes with tumor suppressor roles (*PTPRG*, *XRCC4*) has been described in individual cases. However, recurrent disruption of specific tumor suppressors has not been described^3,8,9,12,13,15–21^ (Supplementary Table 1 and Table 2).

**Table 2:** Summary of Host and Viral Integration Sites.

| Breakpoints in the Human Genome |  |  |
| --- | --- | --- |
| Chromosome | Number of sites | Genes Associated |
| 1 | 14 | PLA2G4A,RABGAP1L,MCOLN2,CASZ1 (I#1) |
| 2 | 6 | SF3B1 (I#1) |
| 3 | 7 | PTPRG (I#1), ATP11B,EEFSEC,ADAMTS9 (I) |
| 4 | 6 | SRD5A2L2,KCNIP4,RELL1,GRID2,FRG1(I), |
| 5 | 21 | TLX3,XRCC4,ISL1,PRLR,CAMK2A(I), CD74(I) |
| 6 | 15 | IL20RA,GMDS,MAP3K7IP2,CDKAL1 (I#4),MRDS1 (I#6),CDKAL1, RP3-348I23.2 |
| 7 | 5 |  |
| 8 | 6 | MYC |
| 9 | 8 | DENND1A;mir601,INPP5E (I) |
| 10 | 3 |  |
| 11 | 8 | PARVA;TEAD1,LUZP2 (I#10;Tr#1),TTC9C (E#1), AHNAK (I) |
| 12 | 3 | AX747640,BEST3 (I#4;Tr#1),KMT2D (I#34) |
| 13 | 1 | DACH1 (I) |
| 14 | 1 | NOVA1 |
| 15 | 3 | ATPBD4;XP_002343377 |
| 16 | 5 |  |
| 17 | 1 | DCAF7 (*I#3) |
| 18 | 2 | CDH2 |
| 19 | 4 | EVI5L,ECH1,GRWD1 |
| 20 | 3 | SNTA1,CBFA2T2,BCL2L1 (E ), TPX2 (I) |
| Y | 1 |  |
| Total | 123 |  |
| Breakpoints in theViral Genome |  |  |
| Genomic Region | Number of sites | Gene |
| Large T Antigen | 42 | LT |
| Small T Antigen | 4 | ST |
| NCCR | 8 |  |
| Viral Capsid Protein 1 | 23 | VP1 |
| Viral Capsid Protein 2 | 10 | VP2 |
| Between LT and VP1 | 2 |  |
| Total | 89 |  |

In brackets : I stands for Intron, Tr is for Transcript and E for Exon. The number following # is the corresponding number of Intron/Transcript/Exon of gene

**Table 3:**
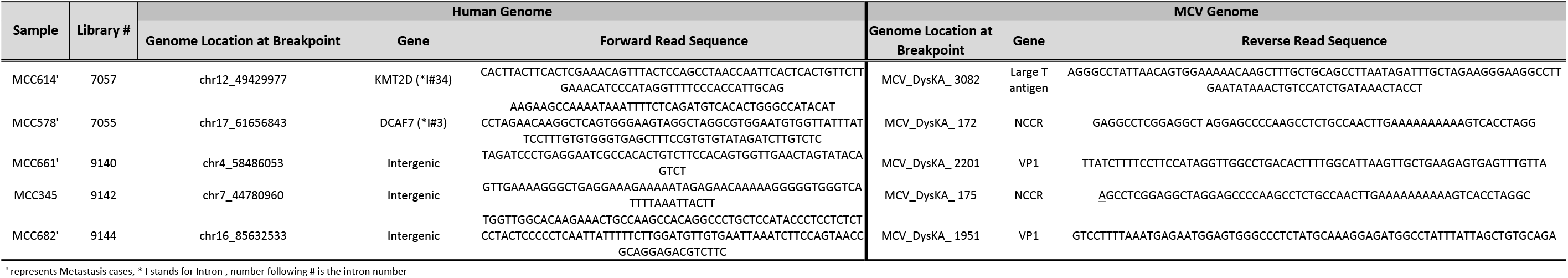
Chimeric Sequences and Novel Integration Breakpoints From Current Study.

Although most tumours demonstrate a single unique genomic integration site, rare exceptions have been described. Starett et al. 2020 found that two independent tumours had overlapping (although non-identical) integration sites on chromosome 1, representing a rare example of a recurrent integration site. An additional case had three distinct integration sites on different chromosomes within a single tumour: one full copy of the viral genome, one smaller segment predicted to retain oncogenic activity (containing NCCR and T antigen coding regions), and one smaller segment predicted to lack oncogenic activity^9^.

Patterns by which the circular MCV viral genome linearizes and integrates have been defined by several studies (Table 1)^8,9^. For both MCV- and HPV-mediated tumours, integration may be followed by the formation of a transiently circular DNA intermediate containing viral genome and flanking regions of the host genome, that can be further amplified through aberrant firing of the viral origin of replication. This piece then reintegrates into the host genome, appearing as amplified regions of the host genome in a tandem head-to-tail conformation interspersed with the viral genome^8,9^.

Although the breakpoint in the MCV genome is highly variable, most studies describe that junctions are predominantly located in exon 2 of the LT gene after the pRb binding site (Table 2). This could result in a truncated, replication-deficient LT protein, equivalent to tumor-specific LT truncating mutations.

More comprehensive characterization of MCV integration locations--and thereby preferences—will advance understanding of virus-mediated tumorigenesis and may reveal therapeutic vulnerabilities. Our study aimed at reviewing published MCV integration site data, and expanding this repertoire of known MCV integration sites. We then adapted a bioinformatics tool designed for HPV integration site detection (HPV Detector/Cancervirus Detector^22^) to detect MCV from previously described MCC whole exome sequencing datasets from patient tumour samples classified as MCV-positive by quantitative PCR^7^. Our results identify additional specific viral and host junction sites, including integration into a tumor suppressor gene frequently mutated in MCV-negative MCC (*KMT2D*). Furthermore, we demonstrate the successful use of HPV Detector/Cancervirus Detector, a simple open-access analysis tool with potential utility in research and clinical testing, for detection of MCV integration in standard whole-exome sequencing data sets.

## RESULTS AND DISCUSSION

### Integration site detection from exome sequencing data

Of the 7 MCC whole exome sequencing datasets, we predicted viral-host junctions in 5 (including 1 primary tumor and 4 metastases). Similar to previous studies, we found that the integration site was highly variable, and at either interchromosomal locations or in introns^8,12,20^. Three of these 5 sites were found in intergenic regions in chromosomes 4, 7 and 16, whereas for two tumors the sites were found in introns of two genes – *KMT2D* (Intron 34) on chromosome 12, and *DCAF7* (Intron 3) on chromosome 17 (Table 1 and Figure 1A). *KMT2D* is a lysine methyltransferase and tumor suppressor gene. *DCAF7* (DDB1 And CUL4 Associated Factor) encodes a scaffold protein for kinase signaling. Unlike a previous report^12^, we did not observe a trend toward involvement of sensory genes.

**Figure 1.**
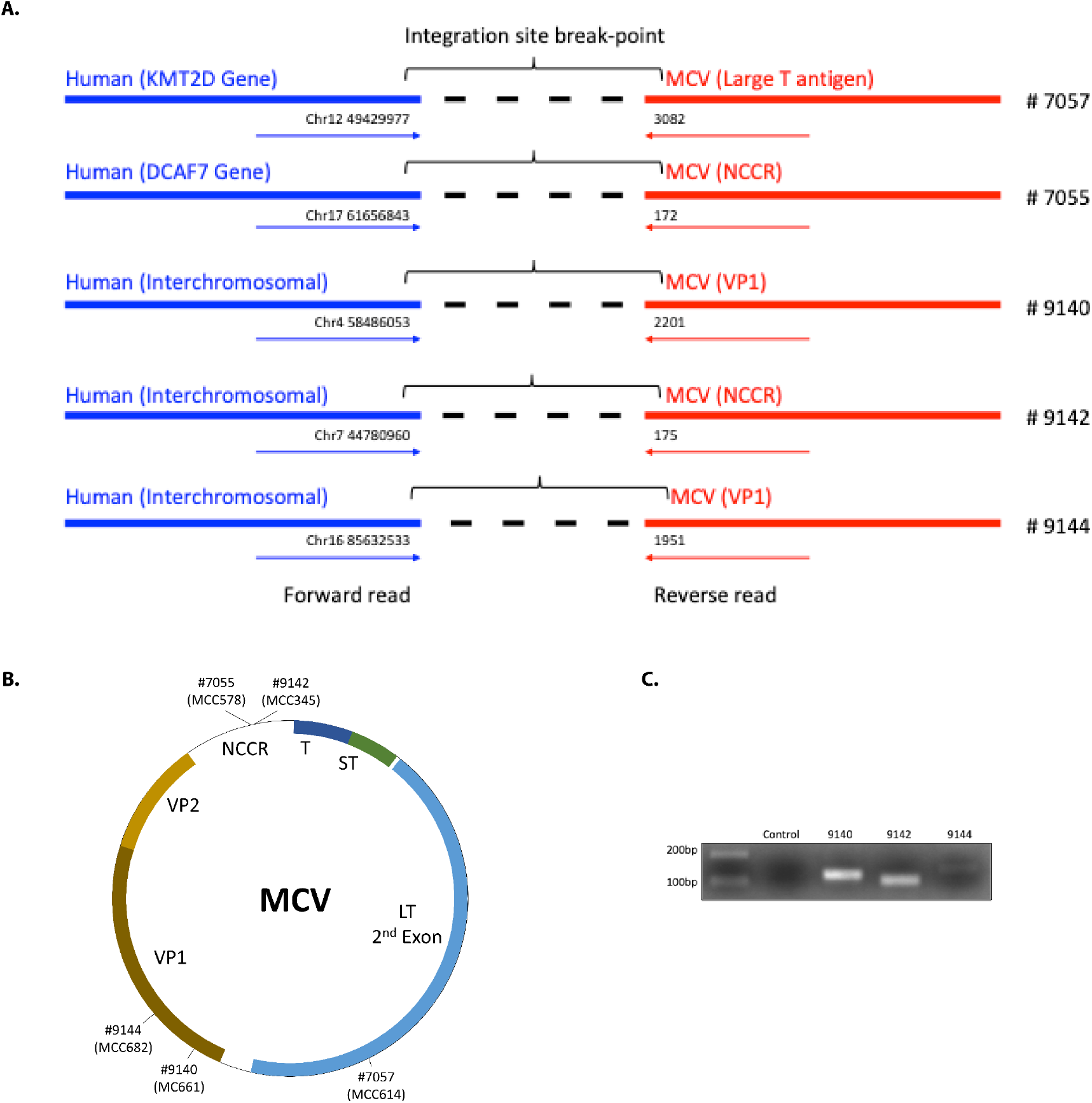
**A.**) Integration breakpoints for the different MCC cases. Blue is the human sequence detected and red is the MCV sequence. **B.**) Integration junctions identified for samples from the current study, by sample number and original tumor ID. **C.**) PCR validation of integration break-points. LT: large T antigen, 2^nd^ exon. NCCR: non-coding control region. ST: small T antigen. T: common T antigen exon. VP: viral protein (capsid).

**Figure 2.**
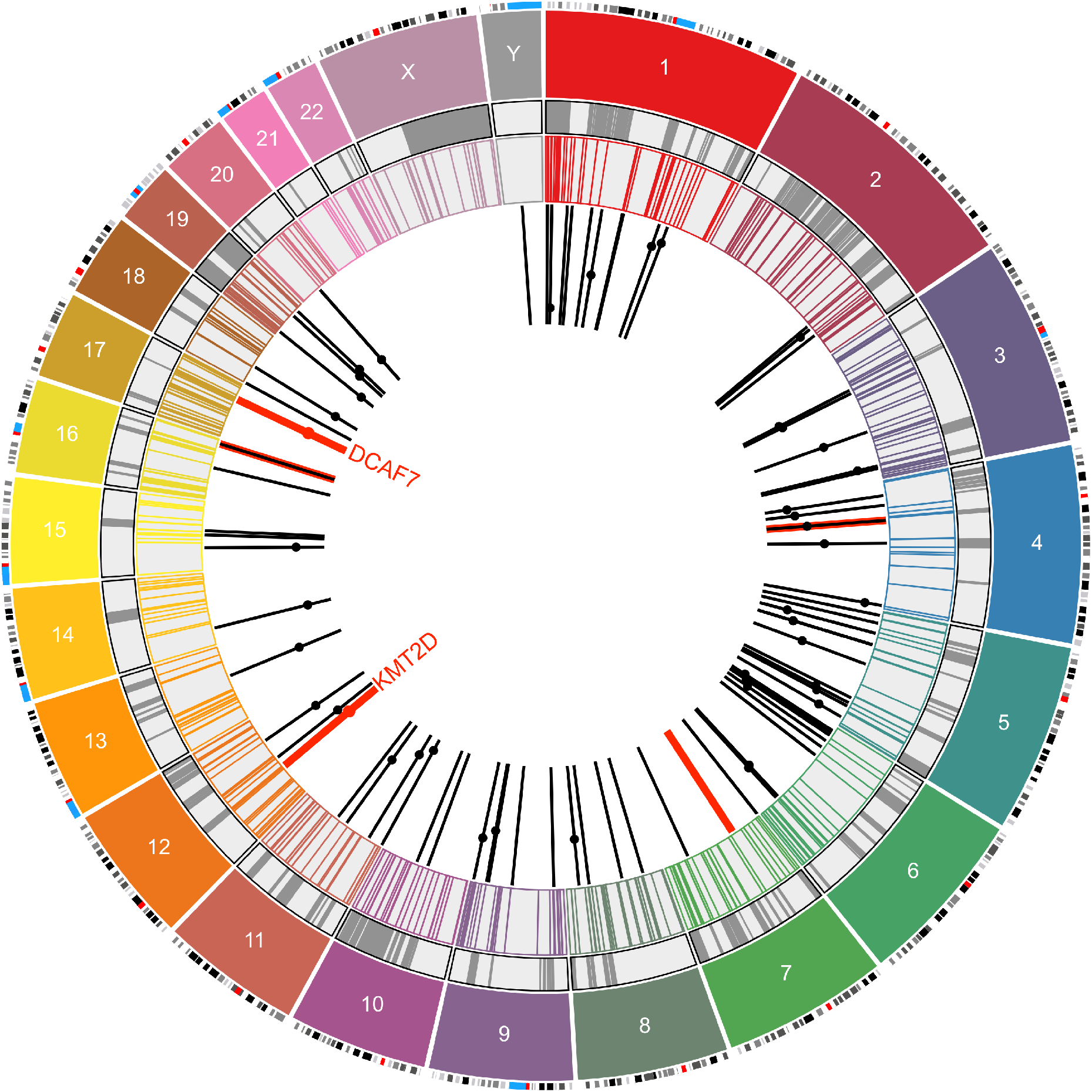
Circo plot representing 96 integration sites of MCV mapped to different chromosomes. The outermost layers represent the chromosomes (colour coded) and their respective chromosome bands. The grey circle marks the human fragile sites from the HumCFS database and the next concentric circle marks the COSMIC database gene sites. The black bars mark 91 integration locations of MCV in MCC from previous reports. Dots on the bars represent the human genes associated with this integration event, their position on the bar is arbitrary. The red bars and dots are data from 5 MCC cases analyzed using HPV Detector in this study. The Circoplot was drawn using the Circa Software (OMGenomics http://omgenomics.com/circa/).

To determine the integration junction for the viral genome, we mapped the sequences to MCV DysKA isolate^23^ (GenBank ID # KX781279.1). The sites we found differed from the previously reported pattern in which integration junctions predominantly involved the LT 2^nd^ exon. In our cases, we found two integration sites in the NCCR region, two in the VP1 region and only one in LT 2^nd^ exon (Table 1 and Figure 1B). Intriguingly, the sequence we found for LT at the junction is preceded by a TAA sequence in the MCV genome, indicating that this integration could also be associated with the generation of a premature LT stop codon, similar to findings by Schrama et al^15^.

By PCR using primers flanking the virus-host junctions, followed by Sanger sequencing of the resulting PCR amplicon, we validated the integration sites predicted for three of these cases (Figure 1 C). The remaining cases lacked adequate remaining DNA for PCR.

### Correlation with previously reported MCV integration sites

We evaluated our 5 integration site results in the context of 91 other previously reported MCV integration sites into the human genome by using Circo Plot^24^ overlaid with COSMIC gene dataset and fragile sites^25^. Chromosome 5 stands out with the highest (21) MCC cases having MCV integrated. There are no integration sites in chromosomes 21, 22 and X. Only five genes of the many associated with MCV integration, namely *SF3B1, TLX3, CD74*, *MYC* and *KMT2D* are cancer-linked genes listed in the COSMIC database.^26^ Additional genes with potential tumor suppressor function that have been shown to be involved by MCV integration include *PTPRG*, *GRWD1*, and *XRCC4*^3,19,21^. Hence, in a minority of MCV-positive MCC tumors, MCV integration itself may be contributing to oncogenesis in a variable and case-specific manner via disruption of cancer genes. Additionally, although there is some overlay with fragile sites, as previously reported, this effect did not appear significant. Furthermore, the integration junctions in the human genome were observed to be enriched in genes having motifs/binding sites of known oncogenic transcription factors such as LEF1 ^27^, MYB ^28^ and AREB6 (ZEB1) ^29^ along with other pathways (Supplementary Table 2); we speculate that this integration pattern might be related to an increased probability of open chromatin for these gene sets.

For the integration junctions in the viral genome (89 total including this study), we observed the previously reported bias to the T antigen region, with 42 cases displaying junctions in LT, and 4 in sT (Table 2).

### Limitations

We were limited by amount and quality of available MCC tumor DNA for PCR-Sanger validation, as the majority was used for next generation sequencing. Similarly, previously described inverse PCR approaches for demonstrating viral genome concatemerization^15^ were not possible due to the limited material. In addition, our approach relies upon identification of off-target capture or shoulder reads of viral sequences within human whole exome sequencing data, thus generating relatively less sequencing depth than a sequencing approach specifically targeting MCV.

### Concluding remarks

Our analysis adds to the repertoire of MCV integration junction data and systematically collates current information regarding the same. We showed unique genetic sites of integration for both the virus and host, confirming the highly variable nature of MCV integration. For the first time we demonstrate integration involving the tumor suppressor gene *KMT2D*, representing the potential inactivation of a tumor suppressor gene also recurrently inactivated in MCV-negative MCC. We also demonstrate that data from tumor whole-exome sequencing data can be effectively interrogated by the simple, open-access Cancervirus Detector tool to demonstrate viral integration sites, thus representing a powerful and accessible new approach for analyzing existing sequencing datasets.

## MATERIALS AND METHODS

### MCV Integration Analysis

HPV Detector uses computational subtraction approach to identify HPV DNA traces from NGS dataset, as published earlier^22^. We modified HPV Detector to align NGS data to Human and MCV genome. In downstream analysis, we subtract all the NGS reads mapping to only human genome. The remaining reads are either mapping to MCV genome completely or are split reads with part of it mapping to human genome and another part mapping to MCV genome. We identified all split reads and annotated them using nearby human and MCV genome regions through BEDTools ^30^. The source code can be accessed via GitHub (https://github.com/pratikchandrani). The sequencing datasets generated previously in Harms et al., 2015^7^ were used for this study. The integration site junctions/genes were further subjected to pathway analysis using GSEA-MSigDB ^31^ using the default parameters.

### PCR

Primers were designed at the flanking regions (one primer in human side and other in viral side) to amplify the integration breakpoints. PCR was performed on genomic DNA from the MCC tumor samples using Platinum Taq DNA Polymerase (catalog # 10966018, Invitrogen), DMSO spike and 20ng of DNA. Water was used as negative control.

### MCV integration site meta-analysis

Based on the closest chromosome band for each integration junction on the human genome, the coordinates were taken from UCSC Browser, GRCh38/hg38. The chromosome and start position were then plotted by Circo plot to represent the integration sites. COSMIC v91 data was downloaded from Sanger Institute website (https://cancer.sanger.ac.uk/census)^26^ and Human Fragile Sites dataset was downloaded from the HumCFS database (https://webs.iiitd.edu.in/raghava/humcfs/)^25^. The Circoplot was drawn using the Circa Software (OMGenomics http://omgenomics.com/circa/). Of the 123 sites for which chromosome number was available, 96 (including 5 from our study) reported information regarding location on the chromosome. Using chromosome band information, circo plot was drawn for these 96 sites. Dots on the bars represent the location of genes in which integration sites were found.

## ACKNOWLEDGEMENTS

We would like to thank Prof. Sudhir Krishna and Prof. Amit Dutt for their valuable support and discussions. We thank Pankaj Vats for assistance with data retrieval and transfer. RA would also like to thank Zaina, Ziyah, and Karan for their love, strength, and support.

## AUTHOR CONTRIBUTIONS

Conceptualization, R.A.; methodology, R.A. and P.C.; validation, J.C.; formal analysis, R.A. and P.C.; investigation, R.A. and P.C.; resources, RA. And P.C.; writing—original draft preparation, R.A.; writing—review and editing, R.A., P.C., J.C., P.H.; supervision, R.A.; funding acquisition, R.A. All authors have read and agreed to the published version of the manuscript.

---

**Supplementary Table 1:**
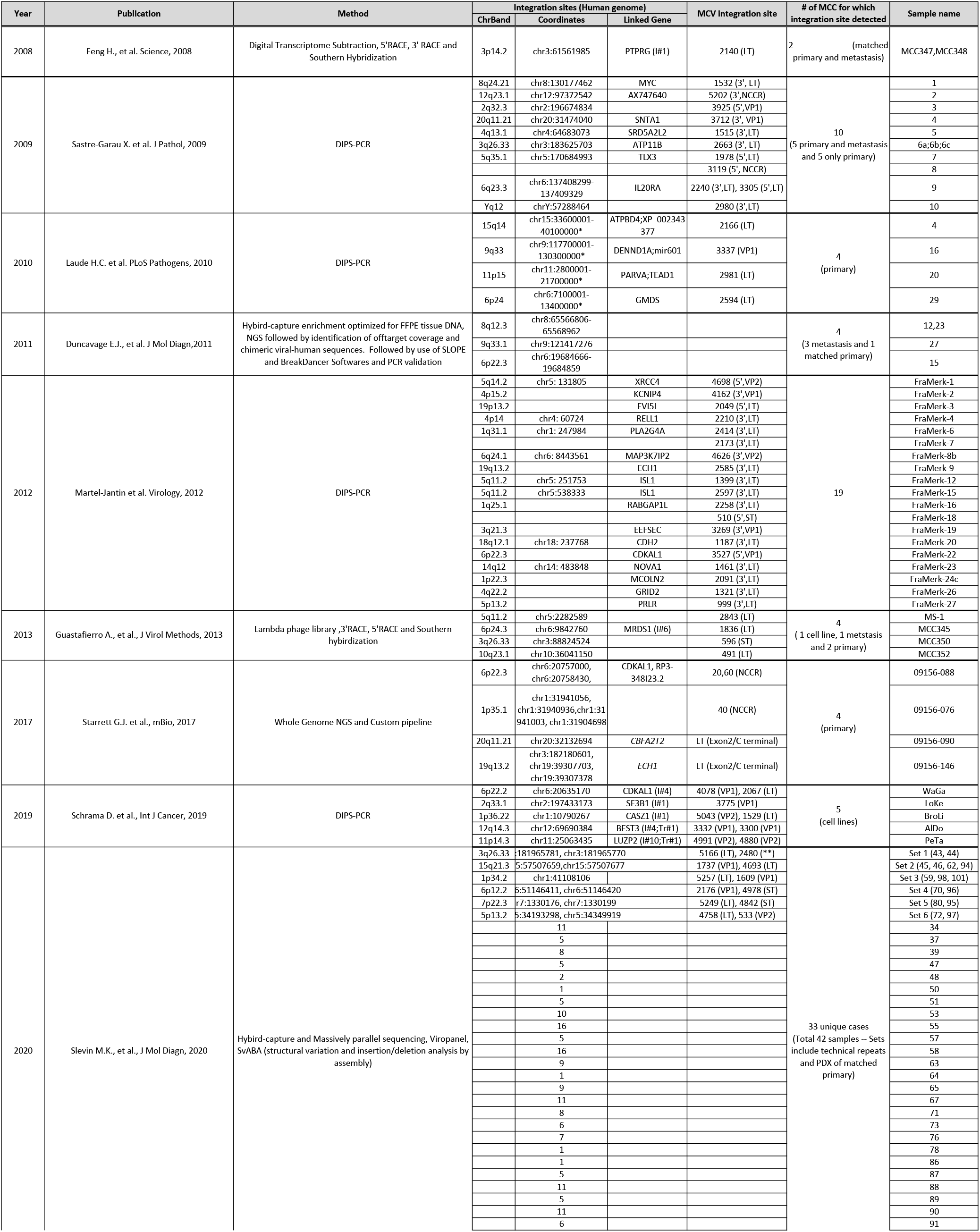
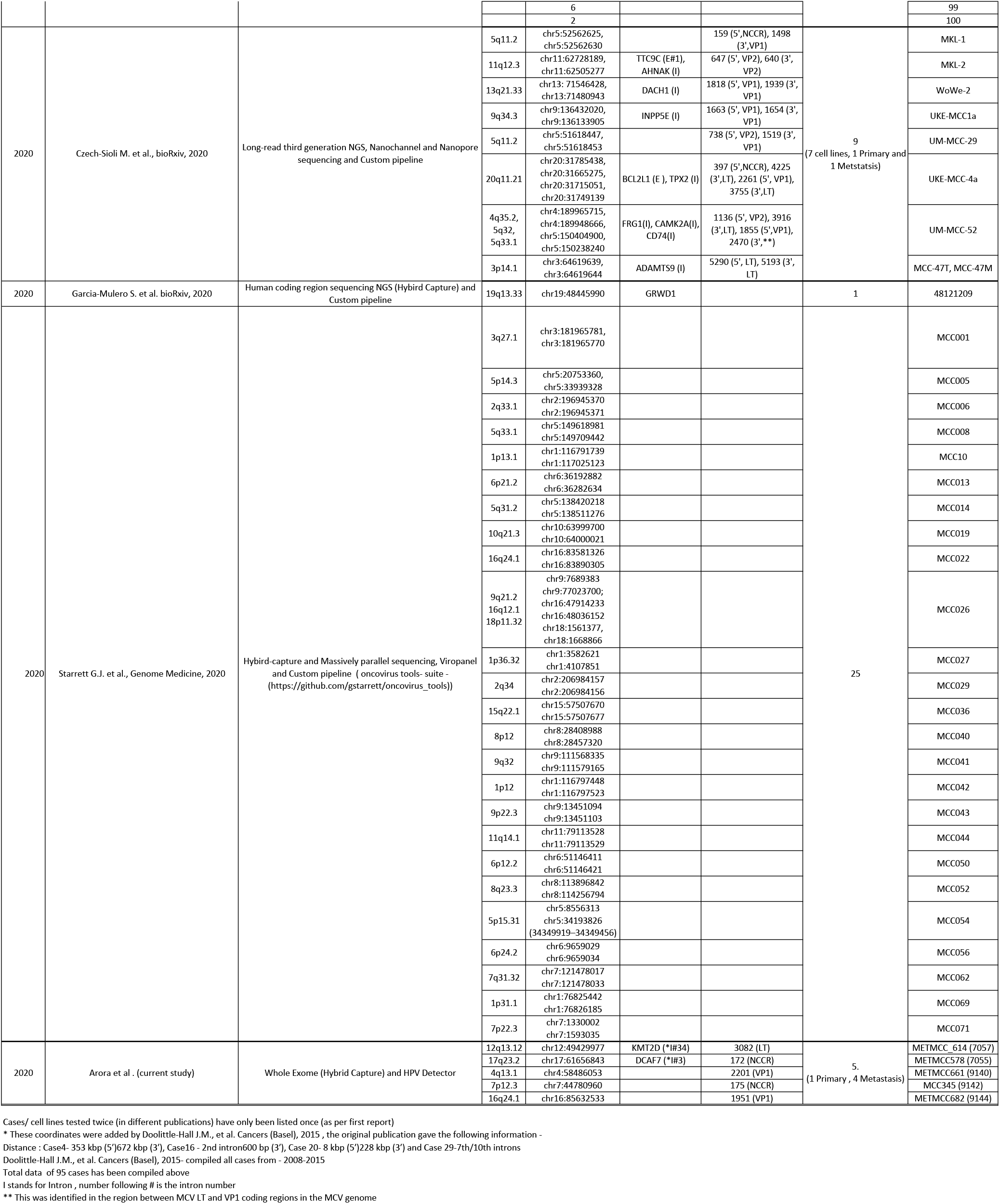
MCV Integration Site Genes and Coordinates From Current Study and Published Reports.

**Supplementary Table 2:**
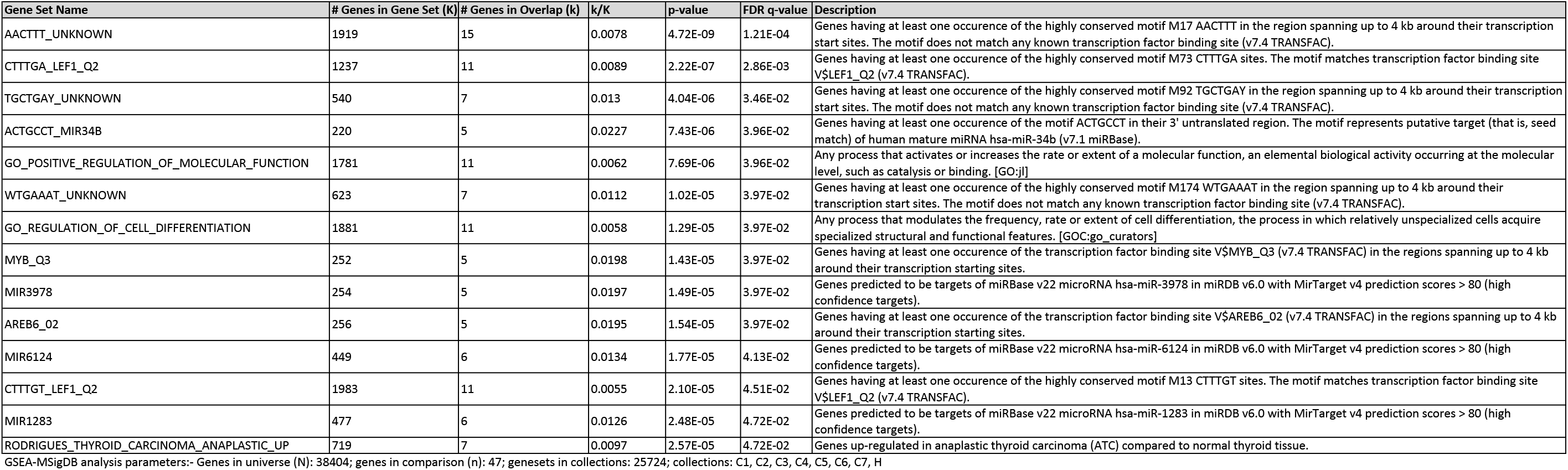
MCV Integration Site Genes and Their Associated Biological Pathways.

